# Anti-infective potential of a quorum modulatory polyherbal extract (*Panchvalkal*) against certain pathogenic bacteria

**DOI:** 10.1101/172056

**Authors:** Pooja Patel, Chinmayi Joshi, Hanmanthrao Palep, Vijay Kothari

## Abstract

Anti-infective potential of a polyherbal ayurvedic formulation namely *panchvalkal* was assayed against three pathogenic bacteria. This formulation was found to exert quorum-modulatory effect on *Chromobacterium violaceum*, *Serratia marcescens*, and *Staphylococcus aureus* at 250-750 μg/mL. Besides altering production of the quorum sensing-regulated pigments in these bacteria, the test formulation also had *in vitro* effect on antibiotic susceptibility, catalase activity and hemolytic potential of the pathogens. *In vivo* assay confirmed the protective effect of this *panchvalkal* formulation on *Caenorhabditis elegans*, when challenged with the pathogenic bacteria. Repeated exposure of *S. aureus* to *panchvalkal* did not induce resistance in this bacterium. To the best of our awareness, this the first report on quorum-modulatory potential of *panchvalkal* formulation, validating the anti-infective potential and moderate prebiotic property of this polyherbal preparation.

## 1. Introduction

Antimicrobial resistance (AMR) is one of the forefront health challenges staring at the mankind in the current times. Modern antibiotic molecules have helped control the infectious diseases to a great extent, however relying solely on these conventional microbicidal antibiotics is not feasible owing to the pathogen’s capacity to evolve to a resistant phenotype [1]. Search for novel alternatives to the conventional antimicrobial therapy is actively being pursued by researchers, and natural products seem to offer a bright ray of hope [2]. In the ancient medicinal texts, there are descriptions of many plant extracts for treatment of microbial infections [3, 4]. Plant products prescribed in ancient literature may be effective, but for their wider acceptance in the modern world, their efficacy has to be demonstrated through appropriate experiments, and also their mode of action needs to be elucidated [5, 6]. All effective antimicrobial formulations may not exert microbicidal effect; they may be exerting their efficacy by targeting virulence of the pathogens without necessarily affecting their growth heavily.

A peep into *Ayurved* (ancient system of Indian traditional medicine) reveals that a *Panchvalkal* formulation (PF) containing the barks of different *Ficus* species has been prescribed as a therapy for maintaining the health of female reproductive system. This formulation has also been mentioned to possess burn- and wound-healing activity [7]. *Charak Samhita* mentions following five plants as the ingredients of PF: *Ficus bengalensis*, *Ficus religiosa*, *Ficus racemosa*, *Ficus lacor*, and *Ficus hispida.* Later *Bhavprakash nighantu* replaced *F. hispida* with *Albezia lebec.* This modified PF has been studied in past for its toxicity, phytopharmacological aspects, and clinical efficacy in the patients suffering from chronic osteomyelitis [8]. We undertook to investigate interaction of this PF with three different pathogenic bacteria viz. *Chromobacterium violaceum*, *Serratia marcescens*, and *Staphylococcus aureus. C. violaceum* is being considered as an emergent pathogen; its infection in humans is reported to be fatal [9, 10]. This bacterium is known to be resistant to penicillins and cephalosporins [11, 12]. *S. marcescens* has also been considered as an emerging opportunistic pathogen causing infections of respiratory tract, urinary tract, meningitis, septicemia, pneumonia, and wound infections [13]. It has been shown to be responsible for frequent nosocomial outbreaks in infants and neonates [14]. *S. aureus* is a major cause of nosocomial infections globally. *S. aureus* infections are often difficult to treat owing to the large population heterogeneity, phenotypic switching, intra-strain diversity, and hypermutability [15]. Pigment production in all these three bacteria is quorum sensing (QS)-regulated. This study assayed the PF for its potential QS-modulatory (QSM) activity against these bacteria, since QS circuit is currently being viewed as a legitimate target for novel anti-virulence agents. QS is an intercellular communication process using acyl homoserine lactone (AHL) or auto-inducing peptides as signal molecules, which regulates expression of a large number of genes including those coding for virulence [16, 17].

## 2. Materials and Methods

### 2.1. Test organisms

*C. violaceum* (MTCC 2656), *S. marcescens* (MTCC 97), *S. aureus* (MTCC 737), and *Lactobacillus plantarum* (MTCC 2621) were procured from MTCC (Microbial Type Culture Collection, Chandigarh), whereas *Bifidobacterium bifidum* (NCDC 255) was procured from NCDC (National Collection of Dairy Cultures), NDRI (National Dairy Research Institute, Karnal). *C. violaceum* and *S. marcescens* were grown in nutrient broth, whereas S. *aureus* was grown in Tryptone Yeast Extract (TYE) broth (ISP HiVeg^™^ Medium No. 1, HiMedia, Mumbai) with additional 0.3%w/v yeast extract. *L. plantarum* was grown in Lactobacillus MRS medium (HiMedia, Mumbai), and *B. bifidum* was grown on MRS with 0.05% Cysteine. Incubation temperature for *C. violaceum*, *S. aureus*, *L. plantarum*, and *B. bifidum* was 37 °C, and for *S. marcescens*, it was 28 °C. Incubation time for *C. violaceum*, *S. marcescens*, *L. plantarum*, and *B. bifidum* was kept 22-24 h, and for *S. aureus*, it was 46-48 h. Antibiotic susceptibility profile of the bacterial strains used in this study was generated using the antibiotic discs-Dodeca Universal – I, Dodeca G - III - Plus and Icosa Universal −2 (HiMedia, Mumbai). *C. violaceum* and *S. marcescens* were found to be resistant to cefadroxil (30 μg), ampicillin (10 μg), cloxacillin (1 μg), penicillin (10 μg). *S. marcescens* showed resistance against vancomycin (30 μg), whereas *S. aureus* was found to be sensitive against all the tested antibiotics.

### 2.2. Test formulation

Capsules of *Panchvalkal* extract (Pentaphyte P5^®^) containing mixtures of bark extracts of *Ficus bengalensis*, *Ficus religiosa*, *Ficus racemosa*, *Ficus lacor*, and *Albizzia lebbec*, were procured from Dr. Palep’s Medical Research Foundation Pvt. Ltd., Mumbai. Powder contained inside the capsules was dissolved in DMSO (Merck, Mumbai) for bioassay.

### 2.3. Broth dilution assay

Assessment of QS-regulated pigment production by test pathogens in presence or absence of the test formulation, was done using broth dilution assay [18]. Organism was challenged with different concentrations (250-1000 μg/mL) of *Panchvalkal* extract. Nutrient broth or TYE was used as a growth medium. Inoculum standardized to 0.5 McFarland turbidity standard was added at 10%v/v, followed by incubation at appropriate temp for each organism. Appropriate vehicle control containing DMSO was also included in the experiment, along with abiotic control (containing extract and growth medium, but no inoculum). Catechin (50 μg/mL; Sigma Aldrich, USA) was used as positive control.

#### 2.3.1. Measurement of bacterial growth and pigment production

At the end of the incubation, bacterial growth was quantified at 764 nm [19]. This was followed by pigment extraction and quantification, as per the method described below for each of the pigment. Purity of each of the extracted pigment was confirmed by running a UV-vis scan (Agilent Cary 60 UV-visible spectrophotometer). Appearance of single major peak (at the *λ*_max_ reported in literature) was taken as indication of purity.

#### 2.3.2. Violacein extraction [20]

One mL of the culture broth was centrifuged (Eppendorf 5417 R) at 15,300 g for 10 min at room temperature, and the resulting supernatant was discarded. The remaining cell pellet was resuspended into 1 mL of DMSO, and vortexed, followed by centrifugation at 15,300 g for 10 min. The purple coloured violacein was extracted in the supernatant; OD was measured at 585 nm. Violacein unit was calculated as OD_585_/OD_764_. This parameter was calculated to nullify the effect of change in cell density on pigment production.

#### 2.3.3. Prodigiosin extraction [21]

One mL of the culture broth was centrifuged at 10,600 g for 10 min. Centrifugation was carried out at 4°C, as prodigiosin is a temperature-sensitive compound. The resulting supernatant was discarded. Remaining cell pellet was resuspended in 1 mL of acidified methanol (4 mL of HCl into 96 mL of methanol; Merck), followed by incubation in dark at room temperature for 30 min. This was followed by centrifugation at 10,600 g for 10 min at 4°C. Prodigiosin was obtained in the resulting supernatant; OD was taken at 535 nm. Prodigiosin unit was calculated as OD_535_/OD_764_.

#### 2.3.4. Staphyloxanthin extraction [22]

One mL of the *S. aureus* culture broth was centrifuged at 15,300 g for 10 min at room temperature. The resulting pellet was suspended in equal volume of methanol, and was kept in water bath (55 C) for 5 min. This was followed by centrifugation at 15,300 g for 10 min, and absorbance of the supernatant containing staphyloxanthin was measured at 450 nm. Staphyloxanthin unit was calculated as OD_450_/OD_764_.

### 2.4. AHL extraction [23]

OD of overnight grown culture of *C. violaceum* was standardized to 1.00 at 764 nm. It was centrifuged at 5000 g for 5 min. Cell free supernatant was filter sterilized using 0.45 μm filter (Axiva, Haryana), and was mixed with equal volume of acidified ethyl acetate [0.1% formic acid (Merck) in Ethyl acetate (Merck)]. Ethyl acetate layer was collected, and evaporated at 55 C, followed by the reconstitution of the dried crystals in 100 μL phosphate buffer saline (pH 6.8). Identity of thus extracted AHL was confirmed by thin-layer chromatography (TLC). R_f_ value of purified AHL, while performing TLC [Methanol (60): Water (40); TLC Silica gel 60 F_254_ plates; Merck] [24] was found to be 0.70, near to that (0.68) reported for N-hexonyl homoserine lactone (C6-HSL) [25].

### 2.5. Antibiotic susceptibility test

After *in vitro* assessment of QSM property of the test formulation, effect of this PF on antibiotic susceptibility of each of the test pathogen was investigated. For this purpose, the most effective quorum modulatory concentration of the PF against a particular bacterium was used. This investigation was done in two different ways. In one set of experiments, we challenged the test pathogen with PF and antibiotic simultaneously, wherein susceptibility of test pathogens to sub-MIC concentration of different antibiotics in absence and presence of test formulation was assessed through broth dilution assay. In another set of experiments the bacterial cells pre-treated with PF were subsequently challenged with antibiotic.

### 2.6. Hemolysis assay [26]

OD of overnight grown culture was standardized to 1.00 at 764 nm. Cell free supernatant was prepared by centrifugation at 15,300 g for 10 min. 10 μL of human blood was incubated with this cell free supernatant for 2 h at 37 °C, followed by centrifugation at 800 g for 15 min. 1% Triton X-100 (CDH, New Delhi) was used as a positive control. Phosphate buffer saline was used as a negative control. OD of the supernatant was read at 540 nm, to quantify the amount of haemoglobin released.

### 2.7. Catalase assay

OD of the culture was adjusted to 1.00 at 764 nm. 400 μL of phosphate buffer was added into a 2 mL vial followed by 400 μL H_2_O_2_. To this 200 μL of the bacterial culture was added, and the mixture was incubated for 10 min. Then 10 μM of sodium azide was added to stop the reaction [27], followed by centrifugation at 12,000 rpm for 10 min. OD of the supernatant was measured at 240 nm to quantify remaining H_2_O_2_ [28], with phosphate buffer as blank.

### 2.8. In vivo assay [29]

*In vivo* efficacy of the test formulation at the concentration(s) found effective during *in vitro* screen was evaluated using the nematode worm *Caenorhabditis elegans* as the model host. Test bacterium was incubated with the PF for 24 h. Following incubation, OD of the culture suspension was equalized to that of the DMSO control. 100 μL of this bacterial suspension was mixed with 900 μL of the M9 buffer containing 10 worms (L3-L4 stage). This experiment was performed in 24-well (sterile, non-treated) polystyrene plates (HiMedia), and incubation was carried out at 22°C. Number of live vs. lead worms was counted everyday till five days by putting the plate (with lid) under light microscope (4X). Standard antibiotic and catechin treated bacterial suspension were used as positive control. Straight worms were considered to be dead. On last day of the experiment, when plates could be opened, their death was also confirmed by touching them with a straight wire, wherein no movement was taken as confirmation of death.

### 2.9. Statistical analysis

All the experiments were performed in triplicate, and measurements are reported as mean ± standard deviation (SD). Statistical significance of the data was evaluated by applying *t*-test using Microsoft Excel^®^. *p* values ≤0.05 were considered to be statistically significant.

## 3. Results

### 3.1. C. violaceum

For *in vitro* studies, *C. violaceum* was challenged with the PF at 250-1000 μg/mL. PF exerted *in vitro* quorum inhibitory effect till 750 μg/mL, but interestingly not at 1000 μg/mL (Figure 1A), whereas growth, though not heavily, was affected at all the test concentrations. Effect of this herbal preparation on growth of *C. violaceum* did not increase with increase in concentration. All the growth inhibition values at different test concentrations ranging from 10.52 - 18.18% did not differ statistically significantly. PF exerted QS inhibitory (QSI) effect at 250-750 μg/mL, with 750 μg/mL being the most effective concentration, causing a reduction of 27.02% in QS-regulated violacein production.

**Figure 1.**
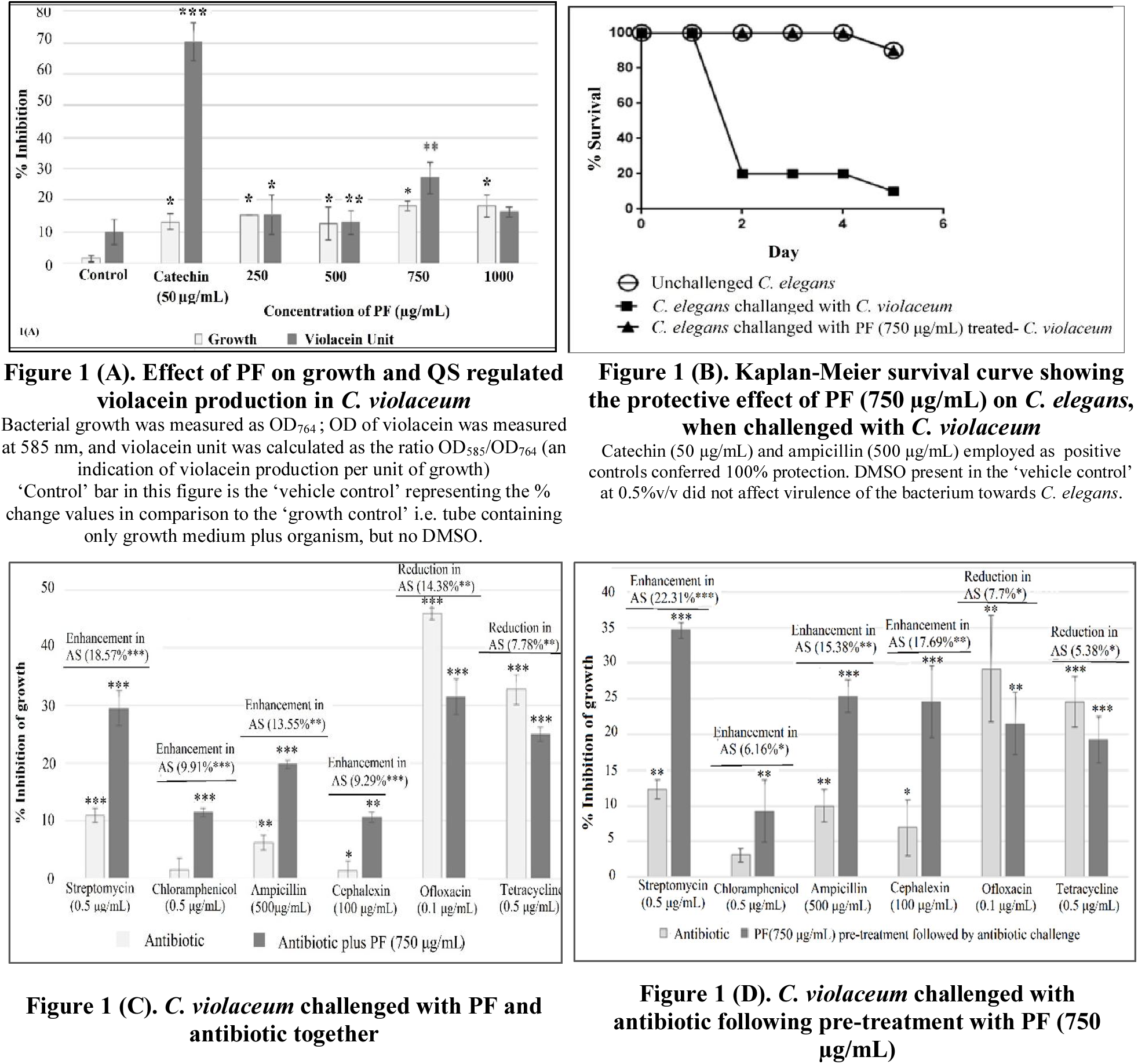
Effect of PF on *C. violaceum*. (^∗^*p*<*0.05*, ^∗∗^*p*<*0.01*, ^∗∗∗^*p*<*0.001*); AS: Antibiotic susceptibility

Once we found 750 μg/mL PF to be the most effective quorum inhibitory concentration against *C. violaceum*, we tried to know whether this PF acts as a *signal supply inhibitor* or *signal response inhibitor.* This was checked by supplementing the QS-inhibited *C. violaceum* culture tube with extracted AHL. The QSI effect of PF was not found to be reversed upon addition of AHL (data not shown), indicating the PF to be acting as a *signal response inhibitor.*

Next we investigated whether the PF affects the bacterial traits other than pigment production i.e. catalase activity, haemolytic activity, and antibiotic susceptibility. Catalase activity of *C. violaceum* culture exposed to PF (750 μg/mL) was enhanced to a minor (2.46%) but significant extent (Table 1), which can be taken as an indication of mild activation of the stress response machinery of the bacterium when challenged with PF. Haemolytic ability, which is an important virulence trait [30], of this bacterium suffered a decrease of 7.01% under the influence of PF (Table 1).

**Table 1:** Effect of *Panckvalkal* formulation on catalase and hemolytic activity of test bacteria.

| Organism | Conc of extract<br>(µg/mL) | % change in<br>catalase activity | % change in<br>Hemolysis |
| --- | --- | --- | --- |
| *C. violaceum* | 750 | 2.46** ± 0.61 | -7.01* ± 2.37 |
| *S. marcescens* | 250 | 1.44* ± 1.17 | -3.95*** ± 1.65 |
|  | 500 | 2.06*** ± 0.37 | -5.42** ± 2.36 |
|  | 750 | 2.13*** ± 0.42 | -5.15*** ± 0.15 |
|  | 1000 | 2.21*** ± 0.71 | -4.65** ± 0.03 |
| *S. aureus* | 250 | 1.82*** ± 0.97 | -4.95* ± 1.60 |
|  | 500 | 2.09** ± 0.76 | -4.42* ± 0.86 |
|  | 750 | 3.59** ± 2.66 | -4.56** ± 0.20 |
|  | 1000 | 4.22*** ± 1.89 | -5.03* ± 1.43 |

\* $p < 0.05$ , \*\* $p < 0.01$ , \*\*\* $p < 0.001$ ; ‘-’ sign indicates a decrease over control; DMSO in ‘vehicle control’ tube had no effect on catalase and haemolytic activity of any of the three bacteria

To investigate whether the susceptibility of *C. violaceum* to different antibiotics is altered under the influence of the QSI formulation, we challenged *C. violaceum* with sub-MIC concentration of six antibiotics belonging to five different classes, in presence as well as absence of PF at its most effective QSI concentration (Figure 1C). Additionally we also tested the effect of antibiotics on *C. violaceum* culture pre-treated with PF i.e. the QS-inhibited culture (Figure 1D).The PF caused an enhancement in the susceptibility of *C. violaceum* to four of the test antibiotics. However, it also caused a reduction in its susceptibility to ofloxacin and tetracycline. These results suggest that though simultaneous use of QS inhibitors and conventional antibiotics may be a good idea, selection of the antibiotic should be done carefully.

After confirming the *in vitro* efficacy of PF on multiple traits (pigment production, catalase activity, haemolysis, and antibiotic susceptibility) of *C. violaceum*, we assayed the PF for its *in vivo* efficacy using the nematode worm *C. elegans* as the model host. The worms were challenged with bacteria previously grown in broth containing (as well as that not containing) 750 μg/mL PF, and their survival was monitored up to five days. Appropriate controls containing the worms challenged with bacteria grown in broth containing 0.5% v/v DMSO or ampicillin (500 μg/mL) or catechin (50 μg/mL) were also included in the experiment. Bacteria previously treated with the PF could kill only 10% of worm population, as against 90% killed by the bacteria not exposed to the PF (Figure 1B). Onset of death in worm population was also delayed by 2 days, when challenged with PF-treated bacteria.

### 3.2. S. marcescens

For *in vitro* studies, *S. marcescens* was challenged with the test formulation at 250-1000 μg/mL. Effect of this PF on *S. marcescens* growth was stimulatory from 250 μg/mL onwards (Figure 2A). Though the effect on prodigiosin production was also stimulatory, magnitude of this effect decreased with increase in concentration. As production of the red pigment prodigiosin in *S. marcescens* is under control of QS [31], the PF can be said to have QSM effect on this bacterium. Catechin used as the positive control exerted stimulatory effect on *S. marcescens* growth, and inhibitory effect on prodigiosin production. As an ideal QS modulator is expected to have minimal or no effect on bacterial growth, we used the PF at 250 μg/mL for further experiments with *S. marcescens*, for the fact that among all evaluated concentrations, this was the concentration exerting lowest effect on bacterial growth, while affecting pigment production maximally.

**Figure 2.**
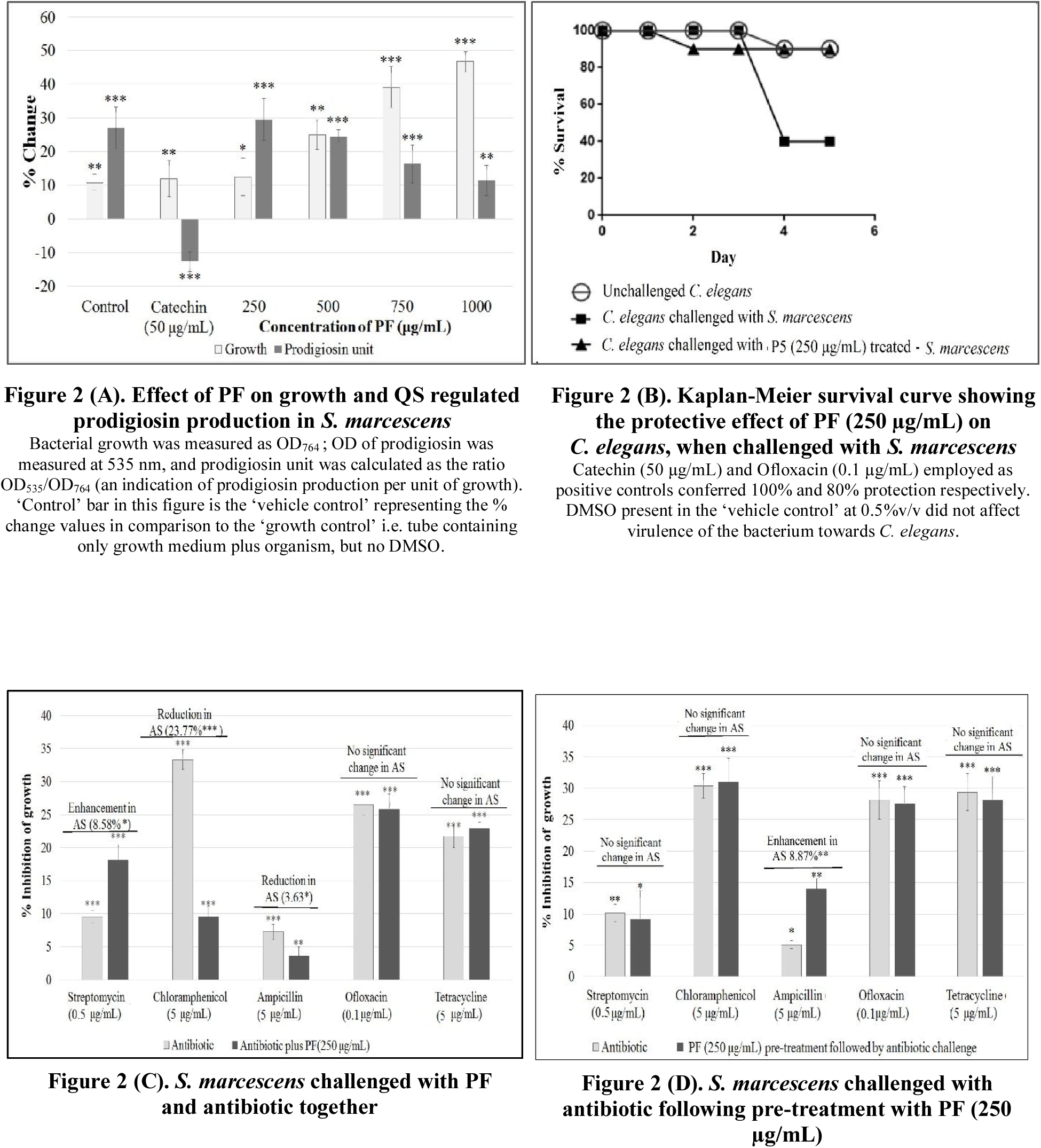
Effect of PF on *S. marcescens*. (^∗^*p*<*0.05*, ^∗∗^*p*<*0.01*, ^∗∗∗^*p*<*0.001*) AS: Antibiotic susceptibility

PF at 250 μg/mL caused a minor (1.44%) but statistically significant enhancement in catalase activity, and reduction in the haemolytic activity (by ~4%) of the *S. marcescens* culture (Table 1). It was also able to alter the susceptibility of this bacterium to three different antibiotics (streptomycin, chloramphenicol, and ampicillin), when tried in combination with these antibiotics (Figure 2C). Susceptibility of this bacterium against chloramphenicol was reduced by nearly 24% in presence of PF; however the *S. marcescens* culture pre-treated with PF did not experienced any alteration in its susceptibility to chloramphenicol (Figure 2D). Antibacterial efficacy of ampicillin in combination with PF was reduced marginally (~4%), but interestingly PF-pre-treated *S. marcescens* became bit more (~9%) susceptible to ampicillin. This might be due to some sort of interaction between ampicillin and the PF, when present together.

*In vivo* assay demonstrated the ability of PF to confer survival benefit on *C. elegans*, *when* challenged with *S. marcescens* (Figure 2B). *C. elegans* challenged with *S. marcescens* receiving no pre-treatment with PF could kill 60 % of worms by 5^th^ day, whereas that treated with PF could kill only 10%. In this case, the survival benefit conferred by PF was equivalent to that conferred by ofloxacin (0.1 μg/mL) used as positive control, as the difference in number of surviving worms for these two cases was statistically not significant.

### 3.3. S. aureus

The PF had a negative effect on the growth of *S. aureus* at all the four test concentrations (Figure 3A), whereas production of the carotenoid staphyloxanthin was enhanced in a dose-dependent fashion. Staphyloxanthin production in *S. aureus* has been known to be regulated by QS [32], and hence the PF can be said to have QSM effect on *S. aureus.* Catalase and haemolytic activity of this bacterium were respectively affected upward and downward, to a marginal extent under the influence of PF (Table 1). Increased catalase activity as well as higher staphyloxanthin production in presence of the PF can be taken as the indication of the bacterium experiencing higher oxidative stress when challenged with this formulation. Carotenoid pigments quench toxic singlet oxygen. They are potent antioxidants making them an important survival factor for detoxifying reactive oxygen species (ROS) [33]. It can be said that the PF used in this study is forcing *S. aureus* to over-activate its antioxidant machinery, by creating oxidative stress for this pathogen.

**Figure 3.**
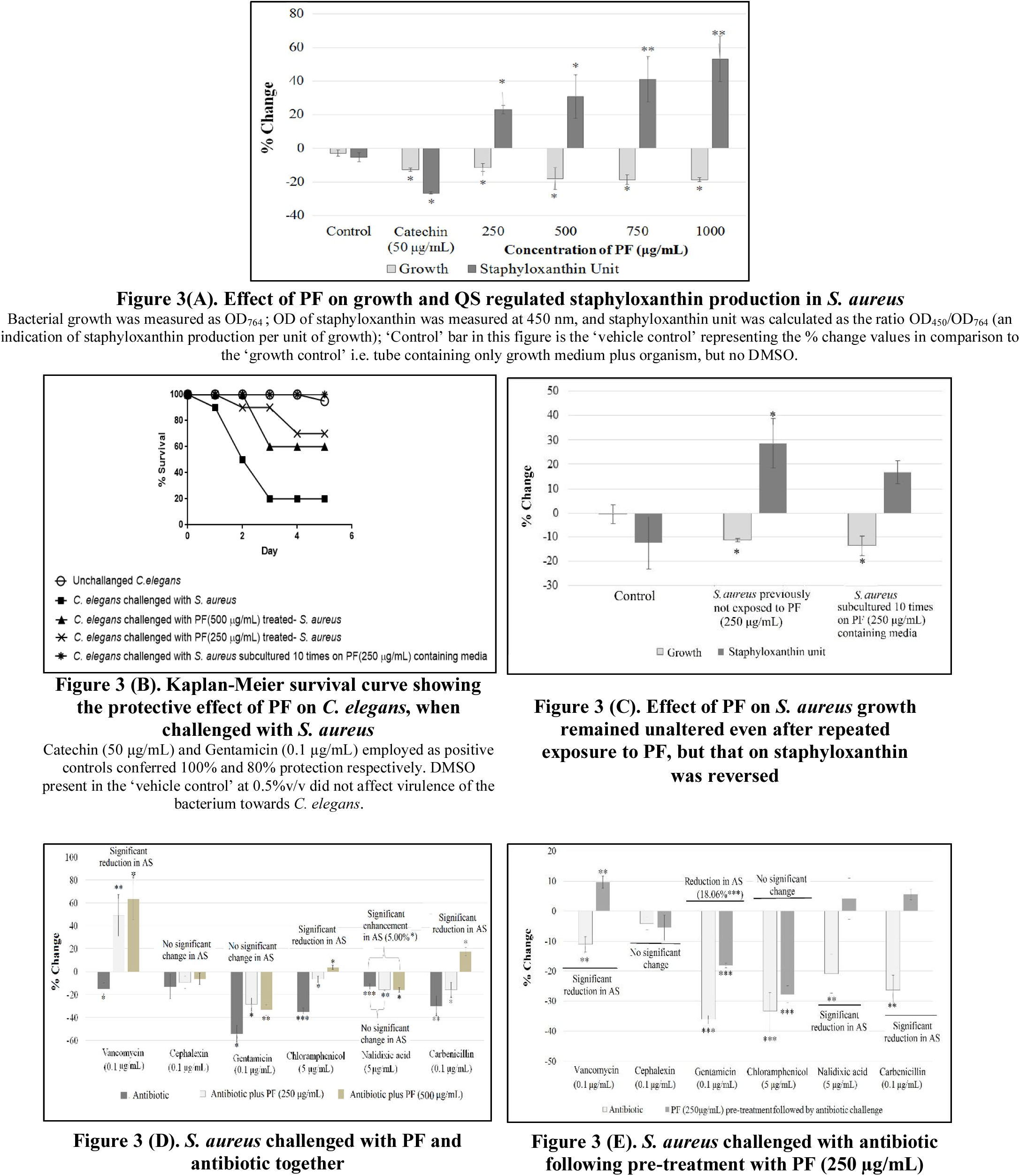
Effect of PF on *S. aureus*. (^∗^*p*<*0.05*, ^∗∗^*p*<*0.01*, ^∗∗∗^*p*<*0.001*) AS: Antibiotic susceptibility antibiotic together

When PF was tried in combination with different antibiotics against *S. aureus*, this bacterium exhibited reduced susceptibility to vancomycin, chloramphenicol, and carbenicillin; and higher susceptibility to nalidixic acid (Figure 3D). PF-pre-treatment resulted in reduced susceptibility towards four of the antibiotics (Figure 3E).

*In vivo* experiments confirmed the anti-infective efficacy of the PF, wherein it offered survival benefit to the nematode worm challenged with *S. aureus.* Interestingly the lower concentration (250 μg/mL) of the PF offered 10% better (*p*<0.001) survival benefit to the worms, than the higher concentration (500 μg/mL) (Figure 3B).

Among the three pathogenic bacteria employed in this study, *S. aureus* is the most dangerous human pathogen. Hence we conducted an additional study with this bacterium to investigate whether repeated exposure of *S. aureus* to PF can induce resistance in this bacterium. For this study, *S. aureus* was subcultured on PF (250 μg/mL) containing agar plates. Culture obtained after such 10 subculturings, was again assayed *in vitro* (Figure 3C) and *in vivo* (Figure 3B). Though the enhancement in staphyloxanthin production observed with ‘control’ *S. aureus* culture (i.e. not previously exposed to PF) was not observed with the PF-pre-exposed culture, the *in vivo* efficacy **remained unaltered**, suggesting that developing resistance against such polyherbal formulations may not be easy for the pathogenic bacteria.

### 3.4. Effect on probiotic strains

An ideal antimicrobial/anti-infective formulation should be selectively effective against pathogenic strains, with no or minimal effect on the human microbiota. We tested the effect of PF on two probiotic strains i.e. *Lactobacillus plantarum* and *Bifidobacterium bifidum* (Table 2). PF had (statistically) similar growth promoting effect on latter at all the concentrations tested. It also promoted growth of *L. plantarum* at 750 μg/mL. Hence it can be said to possess moderate prebiotic and bifidogenic properties.

**Table 2:** Effect of PF on *L. plantarum* and *B. bifidum*.

| Organism | Conc of PF (µg/mL) | % Change compared to control |
| --- | --- | --- |
| *L. plantarum* | 250 | 1.36 ± 1.12 |
|  | 500 | 2.97± 2.73 |
|  | 750 | 8.21*± 3.19 |
| *B. bifidum* | 250 | 18.46* ± 4.76 |
|  | 500 | 20 *± 4.26 |
|  | 750 | 13.84 *± 2.72 |

Bacterial growth was measured as OD<sub>660</sub>; \**p*<0.05

## 4. Discussion

This study has demonstrated the PF to be an effective quorum modulator against gram-positive (*S. aureus*) and gram-negative (*C. violaceum* and *S. marcescens*) bacteria at 250-750 μg/mL. Its spectrum of quorum modulatory action can be said to be broad. Against *C. violaceum*, it seemed to be acting as a *signal*-*response inhibitor*, however it is possible that it being a polyherbal formulation may contain multiple quorum modulatory compounds, some of which may act as *signal*-*supply inhibitor* too. Though modern medicine relies heavily on identification of a single active ingredient to be developed as a therapeutic molecule, ancient traditional medicines have almost always relied on crude extracts from one or multiple plants. Concept of polyherbalism to achieve greater therapeutic efficacy has been mentioned in *Ayurved* too [34]. Whether individual extracts of the bark of constituent plants of PF can exert any meaningful activity remains to be investigated. In context of AMR, the infectious microbes may find it difficult to develop resistance against a multi-component polyherbal formulation, than against a single antibiotic molecule, because in a poly-component formulation there are likely to be multiple compounds exerting antimicrobial action targeting more than one component of bacterial cellular machinery. This may be a reason why the ancient medicinal practices has relied on polyherbal formulations like PF or Ya-Sa-Marn-Phlae (YSMP) for treatment of infections [35].

All the three test bacteria used in this study are involved in a variety of human infections including wound infections. Successful treatment of a wound involves proper healing as well as preventing infections till healing is not complete. PF seems to be capable of enhancing wound healing as well as preventing infection. Wound healing potential of a *Pañcavalkala* formulation in a post-fistulectomy wound was reported by Meena *et al* [36]. Bhat *et al.* reported that PF facilitates wound-healing by reducing microbial load of the wound [37]. However, one of the ingredient plant species mentioned in these studies was different from the PF used in our study. PF has been reported to be useful in treatment of burn wounds and vaginal infections [38, 39].

In the present study, antibiotic susceptibility of the test bacteria was found to alter (increase or decrease) either when antibiotics were used in combination of PF, or against bacteria pre-treated with PF. Pre-treatment with PF could enhance the susceptibility of both *S. maracescens* and *C. violaceum* to ampicillin. Against these two bacteria, streptomycin exerted better efficacy, when used simultaneously with PF. Chloramphenicol exerted better effect against *S. aureus* and *C. violaceum*, but inferior effect against *S. marcescens*, when used in combination with PF. Direction of such modulating effect may be determined by nature of antibiotic applied. In combination with ciprofloxacin and ampicillin, Samoilova *et al.* found certain extracts to provide protective effect, whereas with kanamycin the bactericidal action was enhanced [40]. Mechanisms through which plant products may alter antibiotic susceptibility in bacteria may possibly involve antioxidant, iron-chelating, or prooxidant properties of plant metabolites like polyphenols. Polyphenols were indicated to be involved in conferring protection on *Escherichia coli* against antibiotic toxicity [41].

One of the most famous plant compounds, curcumin was shown to reduce the antimicrobial activity of ciprofloxacin against *Salmonella typhimurium* and *Salmonella typhi* [42]. Combined effect of plant extracts and antibiotics may originate from perturbation of cell membrane and cell wall by the plant extract, resulting in altered influx of antibiotics into the bacterial cells [43]. Hence the lesson is that though combination therapy involving simultaneous use of plant extracts and conventional antibiotics may be a logical idea, this needs to be practiced with caution. Indiscriminate co-administration of antibiotics with herbal formulations may turn out to be therapeutically wasteful or counter-productive, putting the patient at risk, if right combinations are not selected [44].

Though different types of biological activities have been reported in various *Ficus* species [45-47], to the best of our awareness, this study is the first report on QSM potential of PF. Interestingly in this study, *in vitro* efficacy (in terms of % alteration of QS-regulated pigment production) at any particular concentration of PF was found to be lower than its *in vivo* efficacy (in terms of survival benefit conferred on *C. elegans*). This suggests that while screening any test product for its possible anti-infective potential, relatively lower *in vitro* efficacy should not be considered a discouraging factor, as for a quorum-modulator it may not be necessary to inhibit QS to the fullest extent, for being therapeutically useful. Present study is a good demonstration of use of modern scientific approach for validation of traditional therapeutic formulations. More detailed insights can be developed regarding the mode of action of such plant formulations by investigating their effect on bacterial transcriptome and/or proteome profile.

## ACKNOWLEDGEMENT

Authors thank NERF (Nirma Education and Research Foundation) for financial and infrastructural support. PP, CJ, VK thank Dr. H. S. Palep for suggesting ‘panchvalkal’ as a test formulation, and for making the Pentaphyte P5^®^ capsules available for study, and appreciate his zeal for scientific validation of ‘Ayurved’. Biology division, Sophia College for Women, Mumbai is thanked for providing training to CJ on *C. elegans* handling.

## REFERENCES

[1] O’Neill J. Tackling drug-resistant infections globally: final report and recommendations. The review on antimicrobial resistance. Welcome Trust, HM Government; 2016. Available at: https://amr-review.org/sites/default/files/160518_Final%20paper_with%20cover.pdf

[2] Tillotson GS, Theriault N. New and alternative approaches to tackling antibiotic resistance. F1000Prime Reports 2013; 5: 51. Available at: http://dx.doi.org/10.12703/p5-51.

[3] Silva N, Junior AF. Biological properties of medicinal plants: a review of their antimicrobial activity. J Venom Anim Toxins Incl Trop Dis 2010; 16(3): 402–413.

[4] Dias DA, Urban S, Roessner U. A historical overview of natural products in drug discovery. Metabolites 2012; 2(4): 303–336.

[5] Katiyar C, Kanjilal S, Gupta A, Katiyar S. Drug discovery from plant sources: An integrated approach. Ayu 2012; 33(1): 10-19.

[6] Lahlou M. The Success of natural products in drug discovery. Pharmacol Pharm 2013; 4(3), 17–31.

[7] Vaidya ADB. Reverse pharmacological correlates of ayurvedic drug actions. Indian J Pharmacol 2006; 38(5): 311-315.

[8] Palep H, Kothari V, Patil S. Quorum sensing inhibition: A new antimicrobial mechanism of *Panchavalkal,* an ayurvedic formulation. Bombay Hosp J 2016; 58(2): 198-204.

[9] Swain B, Otta S, Sahu K, Panda K, Rout S. 2014. Urinary tract infection by *Chromobacterium violaceum* J Clinic Diagnos Res 2014; 8(8): DD01-DD02.

[10] Bottieau E, Mukendi D, Kalo JR, Mpanya A, Lutumba P, Barbe B, Chappuis F, Lunguya O, Boelaert M, Jacobs J. Fatal *Chromobacterium violaceum* bacteraemia in rural Bandundu, Democratic Republic of the Congo. New Microbes New Infect 2015; 3: 21–23.

[11] Fantinatti-Garboggini F, de Almeida R, Portillo VA, Barbosa TAP, Trevilato PB, Neto CER, Coelho RT, Silva DW, Bartoleti LA, Hanna ES, Brocchi M, Manfio GP. Drug resistance in *Chromobacterium violaceum*. Genet Mol Res 2004; 3(1): 134-147.

[12] Umadevi S, Kumar S, Stephen S, Joseph NM. *Chromobacterium violaceum:* A potential nosocomial pathogen. Am J Infect Control 2013; 41(4): 386-388.

[13] Annapoorani A, Jabbar AKKA, Musthafa SKS, Pandian SK, Ravi AV. Inhibition of quorum sensing mediated virulence factors production in urinary pathogen *Serratia marcescens* PS1 by marine sponges. Indian J Microbiol 2012; 52(2): 160–166.

[14] Rajput A, Shah U, Chauhan B, Shah P. Serratia—An emerging pathogen in hospital environment. Gujarat Med J 2009; 64: 70-71.

[15] Costa AR, Batistao DWF, Ribas RM, Sousa A, Pereira MO, Botelho CN. 2013. *Staphylococcus aureus* virulence factors and disease. In: Méndez-Vilas A, editor. Microbial pathogens and strategies for combating them: science, technology and education. Formatex; 2013: 702-710.

[16] Stevensa AS, Schusterb M, Rumbaughc KP. Working together for the common good: cell-cell communication in bacteria. J Bacteriol 2012; 194(9): 2131-2141.

[17] Holm A, Vikstrom E. Quorum sensing communication between bacteria and human cells: signals, targets, and functions. Front Plant Sci 2014; 5: 309.

[18] Chaudhari V, Gosai H, Raval S, Kothari V. 2014. Effect of certain natural products and organic solvents on quorum sensing in *Chromobacterium violaceum*. Asian Pac J Trop Med 2014; 7: S204–S211.

[19] Joshi C, Kothari V, Patel P. 2016. Importance of selecting appropriate wavelength, while quantifying growth and production of quorum sensing regulated pigments in bacteria. Recent Pat Biotechnol 2016; 10(2): 145–152.

[20] Choo JH, Rukayadi Y, Hwang JK. Inhibition of bacterial quorum sensing by vanilla extract. Lett Appl Microbiol 2006; 42(6): 637-641.

[21] Pradeep BV, Pradeep FS, Angayarkanni J, Palaniswamy M. Optimization and production of prodigiosin from *Serratia marcescens* MBB05 using various natural substrates. Asian J Pharm Clinic Res 2013; 6(1): 34-41.

[22] Song Y, Liu C, Lin FY, No JH, Hensler M, Liu Y, Jeng W, Low J, Liu GY, Nizet V, Wang Andrew HJ, Oldfield E. Inhibition of Staphyloxanthin virulence factor biosynthesis in *Staphylococcus aureus: In vitro, in vivo*, and crystallographic results. J Med Chem 2009; 52(13): 3869–3880.

[23] Chang CY, Krishnan T, Wang H, Chen Y, Yin WF, Chong YM, Tan LY, Chong TM, Chan KG. Non-antibiotic quorum sensing inhibitors acting against N-acyl homoserine lactone synthase as druggable target. Sci Rep 2014; 4: 7245.

[24] McClean KH, Winson MK, Fish L, Taylor A, Chhabra SR, Camara M, Daykin M, Lamb JH, Swift S, Bycroft BW, Stewart GS, Williams P. Quorum sensing and *Chromobacterium violaceum*: exploitation of violacein production and inhibition for the detection of N-acylhomoserine lactones. Microbiol 1997; 143(12): 3703-3711.

[25] Shaw PD, Ping G, Daly SL, Cha C, Cronan JE, Rinehart KL, Farrand SK. 1997. Detecting and characterizing N-acyl-homoserine lactone signal molecules by thin-layer chromatography. Proc Natl Acad Sci USA 1997; 94: 6036–6041.

[26] Neun BW, Ilinskaya AN, Dobrovolskaia MA. Analysis of hemolytic properties of nanoparticles. NCL method ITA-1 Version 1.2, Nanotechnology Characterization Laboratory, Frederick, MD; 2015.

[27] Iwase T, Tajima A, Sugimoto S, Okuda K, Hironaka I, Kamata Y, et al. A simple assay for measuring catalase activity: a visual approach. Sci Rep 2013; 3: 3081.

[28] Weydert CJ, Cullen JJ. Measurement of superoxide dismutase, catalase and glutathione peroxidase in cultured cells and tissue. Nat Protoc 2009; 5(1): 51–66.

[29] Eng SA, Nathan S. Curcumin rescues *Caenorhabditis elegans* from a *Burkholderia pseudomallei* infection. Front Microbiol 2015; 6: Article 290.

[30] Goebel W, Chakraborty T, Kreft J. Bacterial hemolysins as virulence factors. Antonie Leeuwenhoek 1998; 54(5): 453-463.

[31] Khanafari A, Assadi MM, Fakhr FA. Review of Prodigiosin, Pigmentation in *Serratia marcescens* Online J Biol Sci 2006; 6(1): 1–13.

[32] Koch G, Yepes A, Forstner KU, Wermser C, Stengel ST, Modamio J, et al. Evolution of resistance to a Last-Resort antibiotic in *Staphylococcus aureus* via bacterial competition. Cell 2014; 158(5): 1060–1071.

[33] Gaupp R, Ledala N, Somerville GA. Staphylococcal response to oxidative stress. Front Cell Infect Microbiol 2012; 2: 33.

[34] Parasuraman S, Thing GS, Dhanaraj SA. Polyherbal formulation: Concept of Ayurveda. Pharmacogn Rev 2014; 8(16): 73–80.

[35] Chusri S, Tongrod S, Saising J, Mordmuang A, Limsuwan S, Sanpinit S, et al. Antibacterial and anti-biofilm effects of a polyherbal formula and its constituents against coagulase-negative and -positive staphylococci isolated from bovine mastitis, J Appl Anim Res 2017; 45(1): 364-372.

[36] Meena R, Dudhamal T, Gupta SK, Mahanta V. Wound healing potential of *Pañcavalkala* formulations in a postfistulectomy wound. Anc Sci Life 2015; 35(2): 118–121.

[37] Bhat KS, Vishwesh BN, Sahu M, Shukla VK. A clinical study on the efficacy of Panchavalkala cream in Vrana Shodhana W.S.R to its action on microbial load and wound infection. Ayu 2014; 35(2): 135–140.

[38] Joshi J, Rege, V, Bhat R, Vaidya R, et al. Cervical cytology, vaginal pH and colposcopy as adjuncts to clinical evaluation of *Panchavalkal*, an Ayurvedic preparation, in leucorrhoea. J Cytol 2004; 21: 33-38.

[39] Raut AA, Chorghade MS, Vaidya ADB. Reverse Pharmacology. Innovative Approaches in Drug Discovery, Elsevier; 2017. http://dx.doi.org/10.1016/B978-0-12-801814-9.00004-0

[40] Samoilova Z, Smirnovaa G, Muzykaa N, Oktyabrskya O. Medicinal plant extracts variously modulate susceptibility of *Escherichia coli* to different antibiotics. Microbiol Res 2014; 169(2014): 307 – 313.

[41] Smirnova G, Samoilova Z, Muzyka N, Oktyabrsky O. Influence of plant polyphenols and medicinal plant extracts on antibiotic susceptibility of *Escherichia coli*. J Appl Microbio 2012; 113:192–199.

[42] Marathe SA, Kumar R, Ajitkumar P, Nagaraja V, Chakravortty D. Curcumin reduces the antimicrobial activity of ciprofloxacin against *Salmonella Typhimurium* and *Salmonella Typhi*. J Antimicrob Chemother 2013; 68: 139–152.

[43] Sibanda T, Okoh AI. The challenges of overcoming antibiotic resistance: Plant extracts as potential sources of antimicrobial and resistance modifying agents. Afr J Biotechnol 2007; 6(25): 2886-2896.

[44] Eze EA, Oruche NE, Eze CN. 2013. Interaction of the extracts of three medicinal plants with antibiotics against some antibiotic resistant bacteria. Sci Res Ess 2013; 8(28): 1360-1367.

[45] Khan KY, Khan MJ, Ahmad M, Hussain I, Mazari P, Fazal H, et al. 2011. Hypoglycemic potential of genus Ficus L.: A review of ten years of plant based medicine used to cure diabetes (2000–2010). J Appl Pharm Sci 2011; 01(06): 223-227.

[46] Salem MZM, Salem AZM, Camacho LM, Ali HM. Antimicrobial activities and phytochemical composition of extracts of Ficus species: An over view. Afr J Microbiol Res 2013; 7(33): 4207-4219.

[47] Bhalerao SA, Sharma AS. Ethenomedicinal, phytochemical and pharmacological profile of *Ficus religiosa* Roxb. Int J Curr Microbiol App Sci 2014; 3(11): 528-538.

